# Pluripotency factors regulate the onset of Hox cluster activation in the early embryo

**DOI:** 10.1101/564658

**Authors:** Elena Lopez-Jimenez, Julio Sainz de Aja, Claudio Badia-Careaga, Antonio Barral, Isabel Rollan, Raquel Rouco, Elisa Santos, María Tiana, Jesus Victorino, Hector Sanchez-Iranzo, Rafael D Acemel, Carlos Torroja, Javier Adan, Eduardo Andres-Leon, Jose Luis Gomez-Skarmeta, Giovanna Giovinazzo, Fatima Sanchez-Cabo, Miguel Manzanares

## Abstract

Pluripotent cells are a transient population present in the early mammalian embryo dependent on transcription factors, such as OCT4 and NANOG, which maintain pluripotency while simultaneously suppressing lineage specification. Interestingly, these factors are not exclusive to uncommitted cells, but are also expressed during early phases of differentiation. However, their role in the transition from pluripotency to lineage specification is largely unknown. Using genetic models for controlled *Oct4* or *Nanog* expression during postimplantation stages, we found that pluripotency factors play a dual role in regulating key lineage specifiers, initially repressing their expression and later being required for their proper activation. We show that the HoxB cluster is coordinately regulated in this way by OCT4 binding sites located at the 3’ end of the cluster. Our results show that core pluripotency factors are not limited to maintaining the pre-committed epiblast, but are also necessary for the proper deployment of subsequent developmental programs.

## INTRODUCTION

Pluripotency, the ability of a cell to give raise to derivatives of all embryonic germ layers, occurs in cultured embryonic stem cells and for a brief period during development of the mammalian embryo. A small group of transcription factors, most importantly OCT4, NANOG and SOX2, controls this state both in vivo and in culture by regulating a large battery of downstream target genes (1, 2). During preimplantation stages of mammalian embryos, these factors are expressed in the epiblast of the blastocyst, which shares various molecular features with embryonic stem (ES) cells, among them the expression of the core pluripotency factors. It is assumed that progression from pluripotency towards differentiation requires the downregulation of the core pluripotency factors that would lead to the expression of lineage determination genes and turning on of specific developmental pathways. However, the expression of pluripotency factors beyond the blastocyst stage suggests roles not directly related to pluripotency maintenance (3). *Oct4* (official gene symbol *Pou5f1*) is continuously expressed up to embryonic day (E) 8.5, initially throughout the epiblast and subsequently showing progressive restriction to the posterior part of the embryo (4, 5). *Nanog* is re-expressed at E5.5 but only in the posterior-proximal region where it has been shown to control development of the primordial germ cells (6, 7), and is turned off by E7.5 (8). Loss-of-function approaches to investigating the role of *Oct4* and *Nanog* at these stages have proved difficult because preimplantation lethality precludes analysis of later phenotypes (9, 10). To overcome early lethality, conditional *Oct4* mutants have been analyzed at post-implantation. However, loss of Oct4 at these stages leads to tissue disorganization and proliferation defects at gastrulation, what could obscure potential lineage-specific defects (11, 12). Therefore, we still lack a complete understanding of the roles of pluripotency factors during development, as well as how pluripotency and differentiation programs are coordinated in the embryo.

## RESULTS

### Stage-dependent regulation of developmental genes by pluripotency factors

To overcome the limitations or early embryonic lethality and disruption of gastrulation associated with the loss of function of core pluripotency factors, we used doxycycline (dox)-inducible transgenic mouse models providing controlled *Oct4* or *Nanog* expression in post-implantation embryos (13–15). We chose two different time windows for induction of *Oct4* and *Nanog*: from E4.5 to E7.5 and from E6.5 to E9.5 (Fig. 1A), thus maintaining expression beyond the point when endogenous gene activity is turned off. Robust expression of both transgenes was obtained at E7.5 and E9.5 (Fig. S1A), with higher levels in neural tube and mesoderm (Fig. S1B). Nevertheless, expression levels of *Oct4* or *Nanog* in treated embryos were comparable or even lower to those in E14 or R1 (16) ES cells (Fig. S1C). We analyzed the transcriptomes of embryos from untreated and dox-treated females and compared gene-expression changes between stages and models.

**Fig. 1.**
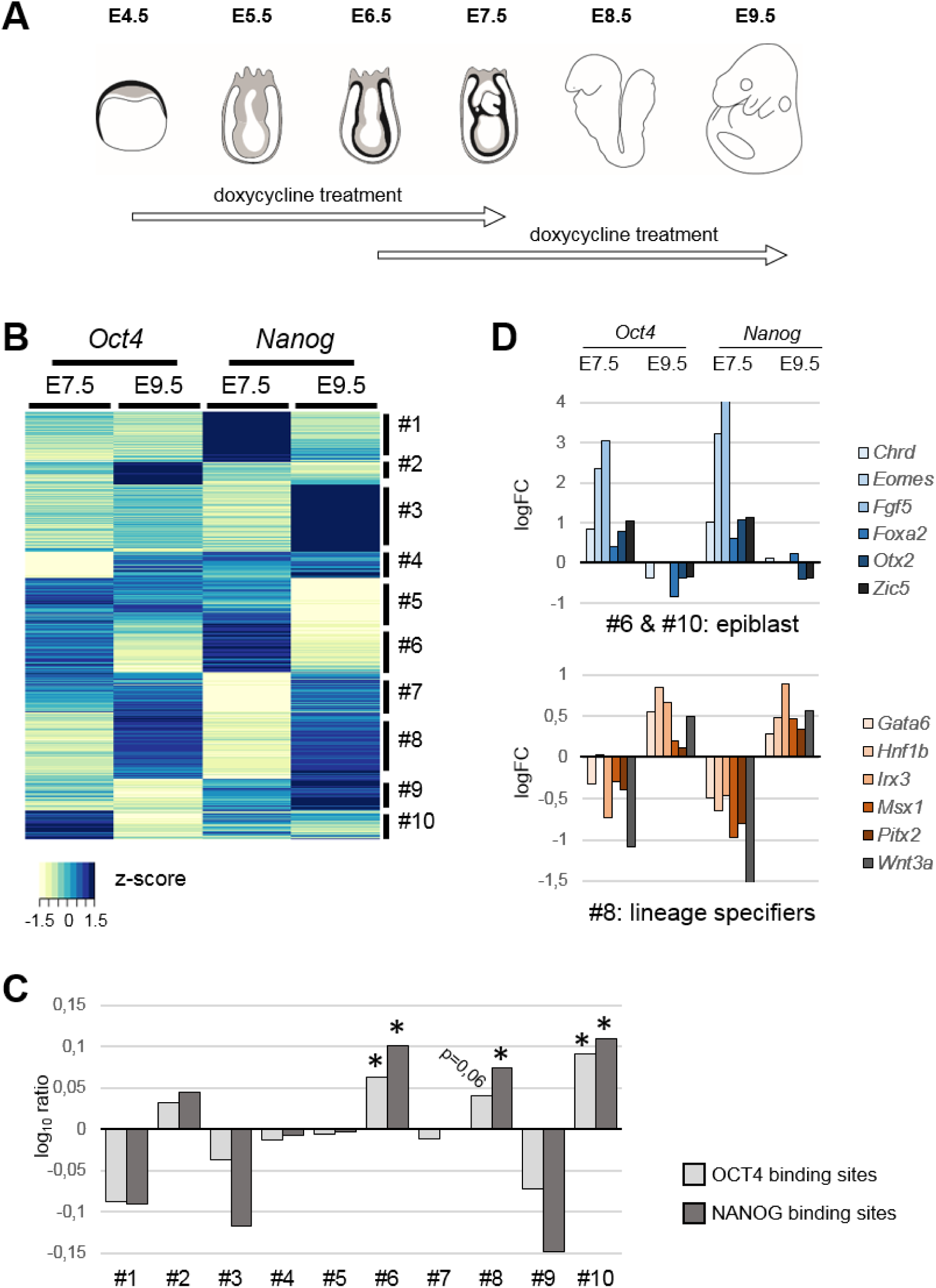
*Oct4* and *Nanog* coordinate developmental programs in the post-gastrulation mouse embryo. (*A*) Diagram showing the windows of doxycycline treatment used to induce *Oct4* or *Nanog* expression at post-implantation stages of mouse development. (*B*) Gene expression changes driven by expression of *Oct4* or *Nanog* for 3 days up to E7.5 or E9.5; unsupervised hierarchical clustering of genes differentially expressed in at least one condition segregated genes into 10 groups according to behavior (clusters #1-#10). (*C*) Enrichment in genes located in the vicinity of ChIP-seq-defined OCT4 or NANOG binding sites, shown as the log10 ratio of each cluster versus genes differentially expressed in at least one of the conditions analyzed. Two-tailed Student *t*-test; +, adjusted *P* value <0.05; *, adjusted *P* value <0.01. (*D*) Representative examples of developmental genes showing opposite behaviors in response to *Oct4* and *Nanog*; genes are either early epiblast markers that are upregulated early and downregulated late (clusters #6 and #10; top panel) or later lineage specifiers that are downregulated early and upregulated late (cluster #8; bottom panel). Expression differences between dox-treated and untreated embryos are shown as logFC of CPMs (counts per million) from RNA-seq data.

More than 50% of genes differentially expressed upon *Oct4* expression up to E7.5 also changed when *Nanog* was induced in the same time window (Fig. S1D), and included major developmental regulators. However, this proportion halved when we compared changes occurring in E9.5 embryos (24%). Similarly, 23% of genes de-regulated by *Oct4* at E7.5 also changed at E9.5. As for *Nanog*, 36% of genes changing at E7.5 were shared with *Oct4*, 16% at E9.5, and only 14% were common at both stages in *Nanog*-expressing embryos (Fig. S1D). The observed degree of overlap between changes induced by *Oct4* and *Nanog* may appear small, in light of their many common target genes in embryonic stem (ES) cells (17); however, this finding is in line with the outcome of overexpressing or knocking-down these factors in culture (18, 19). Core factors of the pluripotency gene regularly network are known to activate each other’s expression (2, 17). We observed positive cross-regulation of *Oct4* and *Nanog* at E7.5, but not at E9.5 (Fig. S1A). Furthermore, we did not observe upregulation of other pluripotency factors, such as *Sox2*, upon *Oct4* or *Nanog* expression. Therefore, there is not an overall activation of the embryonic pluripotency program in the gastrulating embryo driven by these factors.

We performed unsupervised hierarchical clustering of the data (20) using genes that were differentially expressed in at least one condition (4090 genes). Most of the resulting clusters (Fig. 1B) show a stronger tendency for upregulation or downregulation in only one condition (e.g. clusters #1-5; Fig. S2A), confirming the largely independent and stage-specific effects of *Oct4* and *Nanog* expression. Functional annotation showed that most clusters were enriched for genes involved in development and transcription, with some exceptions such as cluster #5, which is enriched for cell-cycle genes, and cluster #7, which includes genes involved in lipid metabolism (Fig. S2B).

To gain insight into functional targets among the de-regulated genes, we checked for the presence of OCT4 or NANOG binding sites in their vicinity by examining published ChIP-seq data in ES cells (21, 22). We assessed for binding near genes from each cluster compared to genes differentially expressed in at least one condition (Fig. 1C). Surprisingly, only three clusters (#6, #8 and #10) showed consistent enrichment for genes located near OCT4 and NANOG sites (Fig. 1C). Moreover, these clusters showed an intriguing pattern in response to *Oct4* and *Nanog*: clusters #6 and #10 were upregulated by both *Oct4* and *Nanog* at E7.5 and downregulated at E9.5, whereas cluster #8 showed the opposite behavior (Fig. S2A). The three clusters are enriched for developmental regulators (Fig. S2B); however, whereas clusters #6 and #10 include early epiblast genes (e.g. *Chordin*, *Dnmt3a/3b*, *Eomes*, *Eras*, *Esrrb*, *Etv5*, *Fgf4*/*5*, *Foxa2*, *Gsc*, *Lefty2*, *Mesp1/2*, *Nodal*, *Otx2*, *Snai1*, *Tdgf1, Zic5*), cluster #8 includes multiple lineage specification genes (e.g. *Cdx1/2/4*, *Dlx5/6*, *Fgf3*, *Gata5/6*, *Gbx2*, *Gli1/2/3*, *Hnf1b*, Hox genes, *Irx3/4/5*, *Meis1/2/3*, *Msx1/2*, *Pitx1/2*, Tbx genes, and *Wnt3a/5a/5b/6*) (Fig. 1D).

### Widespread regulation of Hox genes by pluripotency factors

The large number of Hox genes in this last group (18 genes) prompted us to examine the response of all 39 Hox genes to *Oct4* and *Nanog* (Fig. 2A). *Nanog* has a mild effect on Hox cluster expression (Fig. 2A), mostly downregulating their expression at early stages (14 genes downregulated and 3 upregulated at E7.5, and 5 upregulated at E9.5). On the other hand, *Oct4* has a dramatic effect, significantly downregulating 23 Hox genes when expressed up to E7.5, and upregulating 24 when expressed up to E9.5 (Fig. 2A). It is noteworthy that in this case no Hox gene was upregulated at E7.5 or downregulated at E9.5. *Oct4* affects expression of Hox genes from all four clusters (HoxA-D), but excluding most of the posterior Hox genes from paralog groups 10-13. Interestingly, recent work has shown that *Oct4* delays the activation of posterior Hox genes in trunk progenitors at later embryonic stages (23), suggesting that *Oct4* could have opposite roles in the regulation of anterior and posterior Hox genes during development (24).

**Fig. 2.**
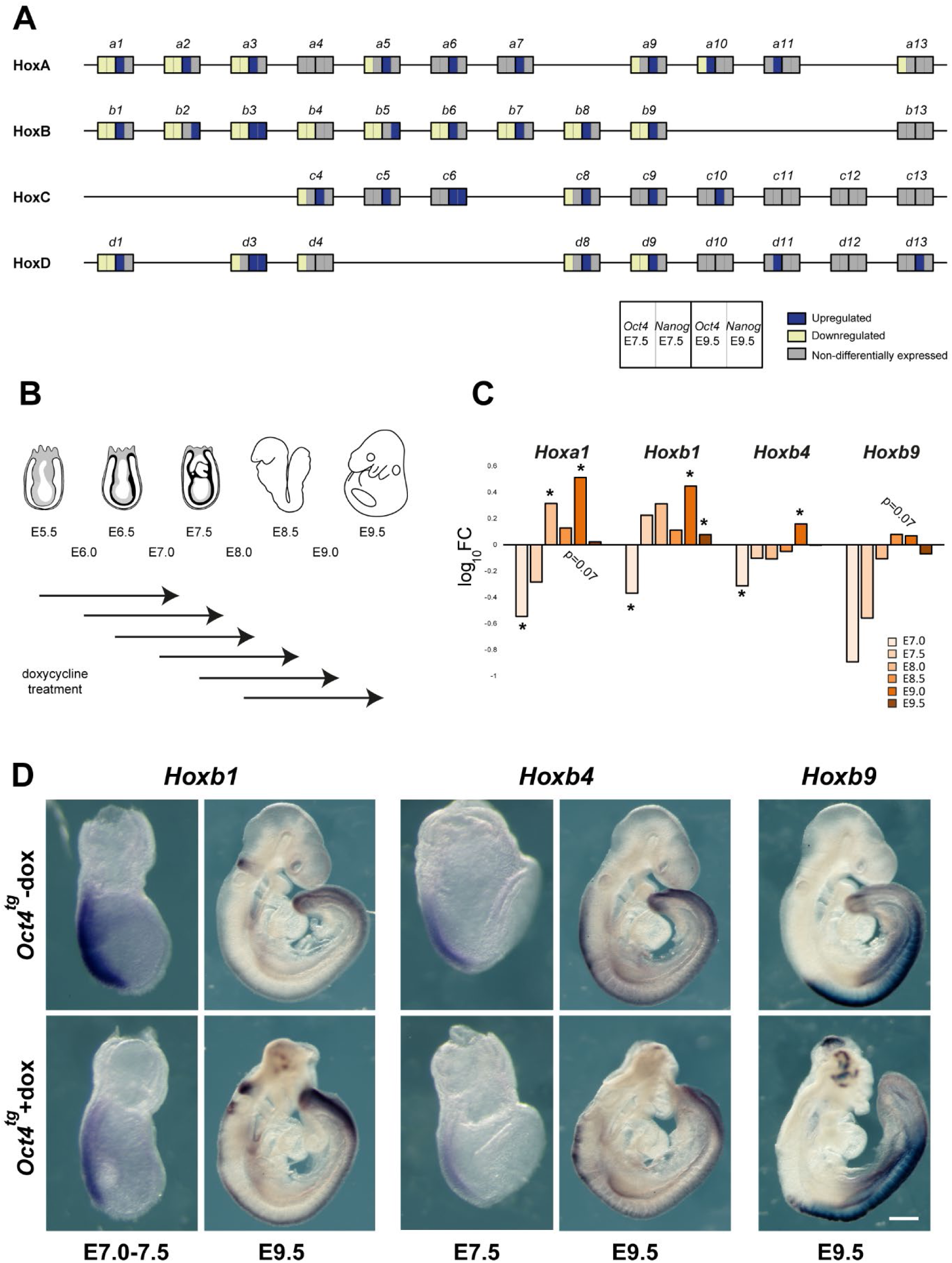
Pluripotency factors mediate a global switch from Hox gene repression to activation. (*A*) Hox cluster gene expression changes induced by *Oct4* or *Nanog* at E7.5 or E9.5, as identified in the RNA-seq analysis. Yellow, downregulated; blue, upregulated; grey, unchanged. (*B*) Diagram depicting the nested windows of 1.5 day dox treatment for *Oct4* induction between E5.5 and E9.5. (*C*) Changes in the expression of selected Hox genes (*Hoxa1*, *Hoxb1*, *Hoxb4* and *Hoxb9*) induced by *Oct4* during development, shown as log ratios of RT-qPCR-measured relative RNA levels in untreated and dox-treated embryos as in B; * *P* value < 0.05 by two-tailed Student *t*-test. (*D*) In situ hybridization analysis of HoxB cluster gene expression in E7.5 and E9.5 untreated (control, top row) and dox-treated (bottom row) *Oct4^tg^* embryos. Scale bar, 140 µm (E7.0-7.5), 170 µm (E7.5), 380 µm (E9.5).

These results strongly suggest that endogenous *Oct4* might be regulating Hox genes during post-implantation development. To address if *Oct4* and Hox genes overlap in the embryo we analyzed published single-cell expression data in E6.5-E7.75 embryos (25) finding that the majority of cells expressing Hox genes co-express *Oct4* (Fig. S3A). This does not rule out the possibility that expression of Hox genes is only occurring in cells that might be turning off *Oct4*, showing low levels of its expression, as a prerequisite for proper lineage commitment. However, Hox expressing cells from E6.5-E7.75 embryos (25) express *Oct4* in an equivalent range to that of all *Oct4* positive cells from the same stages (Fig. S3B). Together, this evidence is compatible with a direct role of *Oct4* in the regulation of Hox clusters.

To determine when the switch from Hox gene repression to activation takes place, we induced *Oct4* for 1.5-day periods between E5.5 and E8.0, harvesting embryos between E7.0 and E9.5 (Fig. 2B). We examined the expression of *Hoxa1* and three HoxB cluster genes (*Hoxb1*, *Hoxb4* and *Hoxb9*) by RT-qPCR using genotype-matched embryos from non-dox-treated females as controls. Consistent with the transcriptomic data, we observed a switch from repression to activation. However, the switch occurred earlier for *Hoxa1* and *Hoxb1* than for *Hoxb4* and *Hoxb9* (Fig. 2C), resembling the collinear activation of the clusters in the early embryo (26, 27).

We next examined *Oct4*-induced changes to HoxB gene expression patterns by in situ hybridization of E7.0-7.5 and E9.5 embryos exposed to dox for 3 days. At E7.0-7.5, expression of *Hoxb1* and *Hoxb4* was downregulated in the posterior of the embryo, as predicted from the transcriptomic analysis, and at E9.5 we observed gain of expression for HoxB genes in several domains of the embryo (Fig. 2D). *Hoxb1* was the most notably affected, with a shift along the anterior-posterior axis (Fig. S4A, arrowhead) together with upregulation or persistence of expression in presumptive rhombomere 6 territory (Fig. S4A, asterisk). Expression of *Hoxb4* conserved its anterior limit of expression (Fig. S4A, arrowhead), but showed an irregular and patchy pattern in the neural tube (Fig. S4A, bracket). As for *Hoxb9*, the anterior limit of expression in the neural tube (Fig. S4A, white arrowhead) shifted anteriorly in comparison with the anterior limit in the paraxial mesoderm (Fig. S4A, black arrowhead). Most strikingly, all three genes examined showed patches of expression in the anterior neural tube (Fig. 2D and Fig. S4B), a territory normally devoid of Hox expression at all developmental stages (28). In situ hybridization in adjacent sections (fig. S4C) revealed that these patches correspond to *Oct4* expressing vesicle-like structures, where the anterior marker *Otx2* was lost and Hox genes were expressed in various combinations (Fig. S4D, arrowheads; Fig. S4E, arrows). These results suggest a coordinated response of the HoxB cluster to *Oct4*, leading to its activation in Hox-free domains such as the forebrain.

On the other hand, *Nanog* caused no obvious changes in Hox gene expression in E9.5 embryos except for an anterior expansion of the *Hoxb1* domain in the neural tube, similar to what we observed for *Oct4* (Fig. S4F).

### Global control of the HoxB cluster by OCT4

To investigate how pluripotency factors regulate Hox genes, we examined previously published ChIP-seq binding profiles of OCT4 and NANOG in mouse ES cells (22, 29). We detected almost no binding within the clusters, with the exception of HoxD, but observed prominent peaks at the anterior ends of the clusters (Fig. 3A and Fig. S5A). For example, in the HoxB cluster, both OCT4 and NANOG bind proximally to the *Hoxb1* promoter (P-site) and to a distal region (D-site) approximately 9 kb downstream of the transcriptional start site (Fig. 3A). These sites have been shown to bind OCT4 during ES cell differentiation (30), and are shared sites of OCT4 binding between ES cells and epiblast-like cells (EpiLC) in the transition from naïve to primed pluripotency (16).

**Fig. 3.**
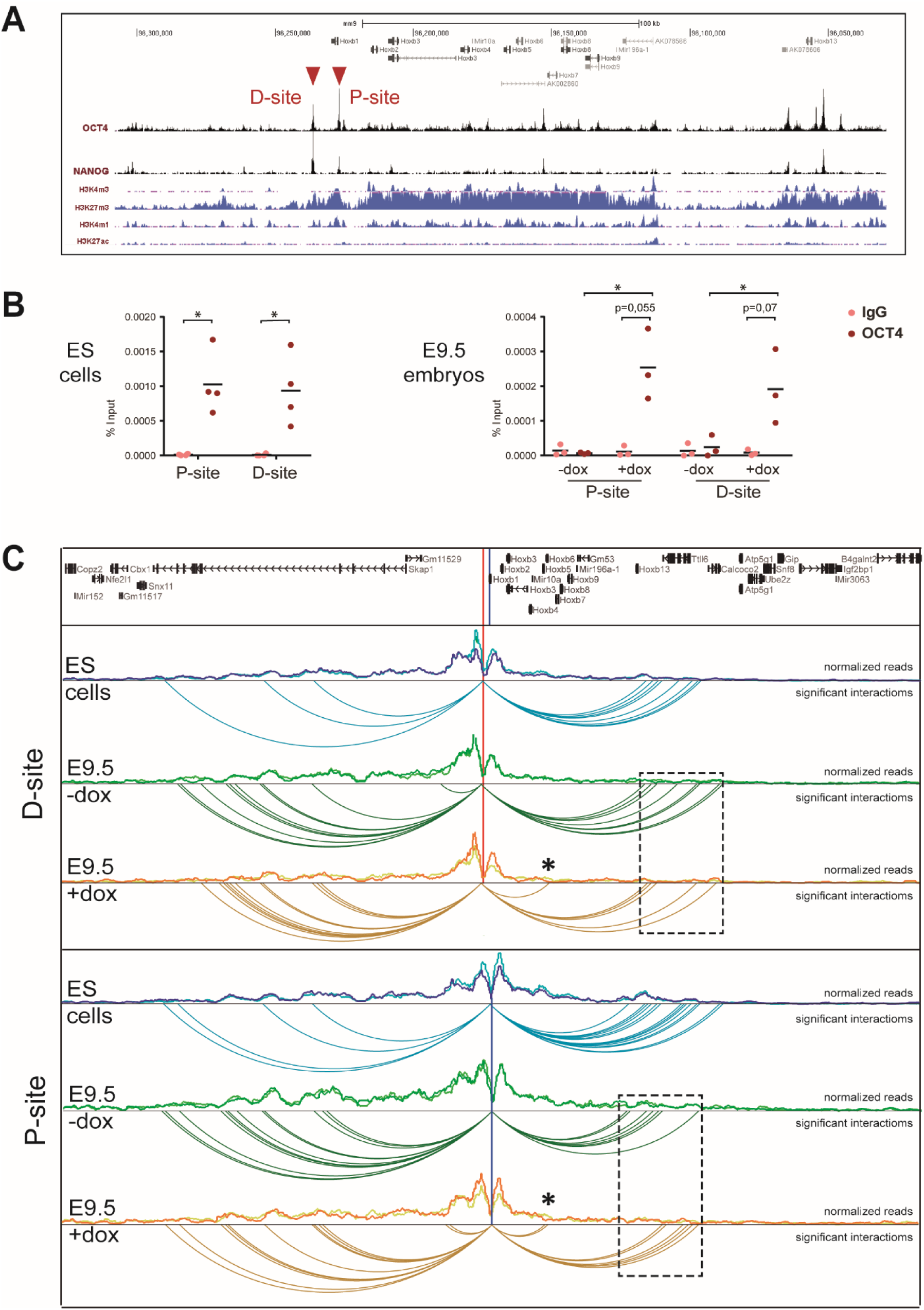
Pluripotency factor binding sites establish long-distance contacts in the HoxB cluster. (*A*) ChIP-seq binding profiles for OCT4 and NANOG (29) along the HoxB cluster (mm9; chr11:96,029,042-96,307,981; reversed). Distribution of histone marks for active transcription (H3K4m3), repression (H3K27m3) and active regulatory elements (H3K4m1, H3K27ac) in Bruce4 ES cells (58) is shown below. Red arrowheads indicate two strong binding sites neighboring *Hoxb1* at the anterior end of the cluster. (*B*) ChIP-qPCR of OCT4 binding to the P- and D-sites in ES cells (left panel, n=4) and in untreated (-dox) and treated (+dox) E9.5 *Oct4^tg^* embryos (right panel, n=3). Enrichment is shown as percentage of input for IgG (negative control) and anti-OCT4 antibody. Horizontal bars indicate means. * *P* value < 0.05 by two-tailed Student *t*-test. (*C*) Chromatin interaction profile established from viewpoints located at the P- and D-sites in a 1 Mb window surrounding the HoxB cluster (mm9; chr11:95,725,993-96,725,993) in ES cells, and untreated (-dox) and treated (+dox) E9.5 *Oct4^tg^* embryos. Distribution of normalized reads in two replicates each is shown, and below significant interactions. Asterisks indicate novel intra-cluster interactions established in dox treated embryos, and boxed regions (dashed line) the interactions established towards the *Hoxb13* region.

We confirmed these sites were occupied by OCT4 in ES cells by ChIP-qPCR (Fig. 3B), using a region from the *Nanog* promoter, previously shown to be bound by OCT4 (29, 31), as a positive control and other unrelated genomic regions as negative controls (Fig. S5B). ChIP in E9.5 embryos showed no binding, but after dox administration we observed binding at both the P- and D-sites (Fig. 3B), demonstrating that these sites are occupied by OCT4 upon induction of its expression.

The response of multiple Hox genes to *Oct4* we observe in the RNAseq dataset and in whole mounts in situs suggests the existence of global mechanisms of Hox cluster regulation by pluripotency factors. To address if these regions could be acting as common regulatory elements for the cluster, we examined their interaction profile by circular chromatin conformation capture followed by high-throughput sequencing (4C-seq) (32, 33). We designed viewpoints for both the P- and D-sites and carried out 4C-seq in mouse ES cells, where OCT4 is present and HoxB genes are not expressed, and in dox treated and untreated E9.5 *Oct4^tg^* embryos (Fig. 3C). Normalized reads were used to fit a distance decreasing monotone function, to take into account that nearby fragments will randomly interact more frequently, and contacts that deviated significantly from the normal distribution were identified.

The chromatin structure surrounding the anterior end of the HoxB cluster is relatively stable independently of its expression (Fig. 3C), as shown for other viewpoints in the cluster (32). Interaction occurs on both sides of the viewpoints: towards the HoxB cluster itself, with a strong limit near *Hoxb13*; and outside of the cluster towards the telomeric region (Fig. 3C). However, interactions are differently distributed around the viewpoint in ES cells and embryos. In ES cells, there are more interactions towards the *Hoxb13* region, possibly reflecting the closed conformation of the cluster mediated by poised promoters (34), whereas in E9.5 embryos interactions increase towards the telomeric gene desert defined by *Skap1* (Fig. 3C), whose expression at E7.0-E7.75 is limited to embryonic blood progenitors (25). This difference in interactions might reflect the active state of the HoxB cluster in the embryo, where distal regulatory elements located in this region (35) are recruited to define its correct expression, as is also the case for the HoxA cluster (36). Our observations are also in line with recent results that show that the HoxA and HoxD clusters are organized in compact domains in ES cells that open up during differentiation (37). Interestingly, when *Oct4* is induced in E9.5 embryos, novel contacts are established from both the P and D sites towards the cluster (asterisks, Fig. 3C) accompanied by a reduction in the interactions with the *Hoxb13* domain (dashed boxes, Fig. 3C). We can conclude that these regions at the telomeric end of the HoxB cluster, which are bound by pluripotency-factors, establish intra-cluster interactions in active or inactive states. Furthermore, the presence of OCT4 in E9.5 embryos leads to a reorganization of the local architecture of the HoxB cluster.

### OCT4 is necessary for proper activation of the HoxB cluster

We next wished to explore the effect of the lack of *Oct4* on the expression of Hox genes. To do so, we analyzed expression of HoxB genes in ES cells depleted of *Oct4*. For this, we used the ZHBTc4 ES cell line that has both copies of endogenous *Oct4* deleted and harbors a tetracycline (tet) dependent *Oct4* transgene, where addition of the drug leads to the quick downregulation of *Oct4* (38). We treated ZHBTc4 ES cells for up to 72 hours (3 days) with tet in 2i culture conditions (39), in order to minimize differentiation of the cells, leading to complete silencing of *Oct4* after the first 24 hours (Fig. S6A). This was accompanied by a more gradual downregulation of *Nanog*, and a sharp upregulation of *Cdx2* as previously described (38, 40) after the first day of tet treatment, that declined on following days (Fig. S6A) possibly as the result of selection against partially differentiated cells in 2i conditions. Genes from the HoxB cluster showed different behaviors in response to loss of *Oct4*. While *Hoxb1* was downregulated in the absence of *Oct4* at all days, *Hoxb4* showed initial upregulation at day 1 but downregulation by day 3. On the other hand, *Hoxb9* was upregulated throughout the 3-day culture (Fig. 4A). Therefore, *Oct4* is necessary for basal activation of 3’ genes, while needed to repress expression of further 5’ HoxB genes.

**Fig 4.**
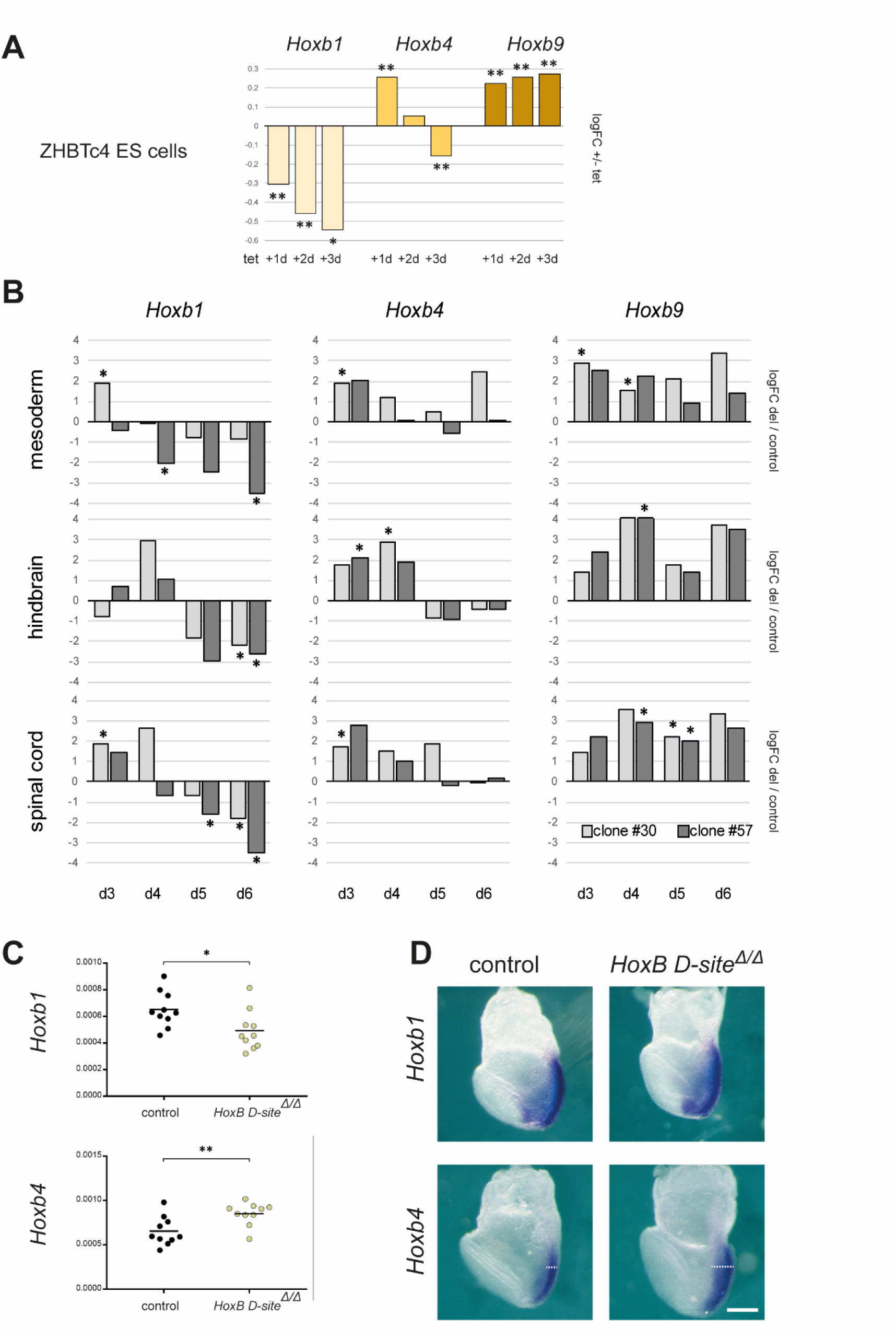
OCT4 is required for correct expression of HoxB genes during differentiation. (*A*) Changes in the expression of *Hoxb1*, *Hoxb4* and *Hoxb9* in ES cells upon *Oct4* depletion. Differences in expression are shown as log-fold-change of RT-qPCR-measured relative RNA levels measured by RT-qPCR in ZHBTc4 ES cells treated with tetracycline for three days (d1-d3) versus untreated cells. Statistical significance was calculated by Fisher’s Least Significant Difference test; n=3; *P* value: * <0.05, ** <0.01. (*B*) Changes in the expression of *Hoxb1*, *Hoxb4* and *Hoxb9* as a consequence of the deletion of the distal OCT4 binding site up to six days of differentiation (d3-d6) of ES cells to mesoderm, hindbrain, or spinal cord lineages. Differences in expression are shown as log-fold-change of RT-qPCR-measured relative RNA levels in two independent deleted homozygous clones (#30, light gray; #57, dark gray) versus control ES cells. n=3; * *P* value < 0.05 by two-tailed Student *t*-test. (*C*) Expression of *Hoxb1* and *Hoxb4* in E7.5 control (blue) and *HoxB D-site^Δ/Δ^* (pale yellow) embryos as quantified by RT-qPCR. n=10; *P* value: * < 0.05, ** < 0.01 by two-tailed Student *t*-test. (*D*) Expression of *Hoxb1* (top) and *Hoxb4* (bottom) in E7.5 early headfold stage *HoxB D-site^Δ/Δ^* (right) embryos compared to controls (left); n=6. Dashed lines indicate the extension of *Hoxb4* expression in the posterior region of the embryo. Scale bar, 200 µm.

### Deletion of the distal OCT4 site disrupts the pattern of HoxB expression during differentiation

To complement these observations, we decided to test the necessity of the OCT4 bound regions described above in the regulation of the HoxB cluster by deletion in ES cells by CRISPR/Cas9 mediated genome editing (41). We examined the genomic regions covered by ChIP-seq peaks, finding that the proximal site (P) contains one consensus OCT4 binding site, while the distal site (D) at least two (Fig. S6B). In the case of the P-site, this consensus lies within the previously described proximal *Hoxb1* auto-regulatory element (42) and in very close proximity to the promoter. Given the difficulty of deleting this site without compromising other known regulatory inputs on *Hoxb1*, we decided to only analyze the effect of deleting the D-site. This region does not map to any other known *Hoxb1* regulatory elements, such as the two described 3’ retinoic acid response elements (43, 44) (Fig. S6B).

We generated two independent ES cell clones deleted for the D site (clone #30 and #57; Fig. S6B) and analyzed changes in HoxB gene expression during their differentiation towards mesodermal or neural fates (Fig. S6C) (45, 46). Comparison of parental with HoxB D-site^Δ/Δ^ ES cells along differentiation to mesodermal, hindbrain (with low dose of retinoic acid) or spinal cord (high dose of retinoic acid) showed a similar trend for each of the Hox genes examined independently of the differentiation protocol, and a comparable behavior in the two independent clones (Fig. 4B). *Hoxb1* is initially expressed at higher levels in deleted cells, but is downregulated as compared to controls at later stages of differentiation. *Hoxb4* also shows initial upregulation, but no significant changes at later time points. Finally, *Hoxb9* is consistently activated in deleted cell lines throughout the differentiation window analyzed (Fig. 4B), as also occurs when *Oct4* is depleted from ES cells (Fig. 4A).

To analyze the effect of the deletion in vivo, we used the HoxB D-site^Δ/Δ^ ES cells to generate mouse lines. Homozygous mice survive to term, what did not come as a surprise, as deletion of the entire HoxB cluster in homozygosity does not cause embryonic lethality (47). We examined the expression of *Hoxb1* and *Hoxb4* at E7.5, a time when anterior HoxB genes and *Oct4* are still co-expressed (Fig. S3A, B). RT-qPCR showed down-regulation of *Hoxb1* and upregulation of *Hoxb4* in deleted embryos (Fig. 4C). This further confirms the different roles of *Oct4* at this stage, when it would be necessary for proper levels of express info *Hoxb1* but still lowering *Hoxb4*.

Whole mount in situ hybridization at the early head-fold stage showed no clear changes in *Hoxb1*, but an expansion of the expression in the primitive streak region at the posterior part of the embryo of *Hoxb4* (white dashed line, Fig. 4D). We also examined expression of HoxB genes in E9.5 embryos from the *HoxB D-site^Δ/Δ^* line, and did not observe mayor changes except for overall lower levels of *Hoxb4* expression (Fig. S6D). Therefore, these results suggest that at early stages of expression, OCT4 is necessary to fine-tune expression of HoxB genes in their endogenous domains.

## DISCUSSION

It is generally assumed that pluripotency factors act to restrict lineage decisions before gastrulation. However, *Oct4* has been shown to participate in several later developmental decisions, including primitive endoderm development (48), lineage priming (30, 49, 50), primitive streak proliferation (12), and the regulation of trunk length (23). Furthermore, recent results have shown how the lack of *Oct4* in early gastrulating embryos results in a blockade of epithelial-to-mesenchymal transition in the posterior epiblast, thus disrupting proper axis formation (11). Both single-cell RNA-seq data and in situ detection show that *Oct4* and Hox genes are co-expressed in cells of the gastrulating embryo, from the onset of Hox gene expression and up to E8.5 (4, 5, 25). We propose that at these early stages, *Oct4* plays a dual role, first maintaining Hox genes silent before lineage commitment, and later being required for their proper activation. This behavior is specific for *Oct4*, as in the case of *Nanog* we only observe initial repression of Hox genes, which agrees with the recent description of the mutual cross-repression of *Nanog* and *Hoxa1* to differentially regulate a common set of downstream target genes involved in early phases of lineage commitment (51).

The concerted response of Hox genes, together with the chromatin interactions established by bound regions we see in the HoxB cluster, suggests that *Oct4* regulates globally Hox clusters and forms part of the complex regulatory apparatus that ensures proper Hox gene expression (28, 36). Furthermore, when we examine the expression of anterior, middle and posterior HoxB genes in gain- and loss-of-function models, we observe a collinear response to *Oct4*, in line with recent findings of the repression of most posterior 5’ Hox genes by *Oct4* (23, 24).

Moreover, our expression data indicate that other patterning genes respond in a similar fashion, suggesting that OCT4 and other pluripotency factors mediate a switch from repression to activation of an array of developmental regulators at the time of lineage decisions. Therefore, initial lineage specification involves not only dismantling of the core pluripotency gene regulatory network (3), but also a switch in function of key factors such as OCT4 from repressors to activators that would supervise the transition from pluripotency to lineage determination.

## MATERIALS AND METHODS

### Animal models

Double-homozygote transgenic males of the *Oct4*/rtTA (R26-M2rtTA;Col1a1-tetO-Oct4) (14) or *Nanog*/rtTA (R26-M2rtTA;Col1a1-tetO-Nanog) (15) mouse lines were mated with CD1 females, which were treated with 0.2 or 1 mg/ml dox in the drinking water to induce the *Oct4* or *Nanog* transgene respectively in embryos. For *Oct4* transgene induction in E7.5 embryos to be used for in situ hybridazation, a single 100μl intraperitoneal injection of 25µg/µl doxycycline was administered to pregnant females at E5.5, followed by 0.5 mg/ml dox administration in drinking water. Non treated mice of the same genotype were used as controls. Mouse lines deleted for the *Oct4* distal site adjacent to *Hoxb1* were generated by blastocyst injection of mutated ES cells (see below) following standard procedures (52).

### RNA-seq

RNA-seq was performed with three biological replicates, each consisting of pools of 8-12 E7.5 embryos or three E9.5 embryos obtained from untreated (controls) or doxycycline-treated double heterozygous embryos. Equally, three biological replicates of E14 ES cells were used for RNA-seq. Single-end sequencing was performed by the CNIC Genomics Unit using a GAIIx sequencer. Adapters were removed with Cutadapt v1.14 and sequences were mapped and quantified using RSEM v1.2.20 to the transcriptome set from Mouse Genome Reference NCBIM37 and Ensembl Gene Build version 67. Differentially expressed genes between groups were identified using the limma bioconductor package. Only p-values < 0.05 adjusted through Benjamini-Hochberg procedure were considered as significant. Clustering analysis was conducted for all genes differentially expressed between the induced and control conditions for any of the four conditions. Overrepresented biological categories were identified using DAVID v6.8 (53). Lists of genes located close to OCT4 and NANOG binding sites were generated from published ChIP-seq datasets (21, 22).

Single-cell gene expression data from Scialdone et al. (2016) was analyzed to assess co-expression of *Oct4* and Hox gene. A gene is considered to be expressed in a cell when its expression level is > 0 FPM (fragments per million).

### RT-qPCR assays

Total RNA from single embryos from E8.0 onwards, embryo pools up to E7.5, or ES cells directly lysed in their wells, was extracted with RNeasy kit (Qiagen) and digested with DNAse I (Qiagen) to remove genomic DNA. 0.5-1.0 μg of total RNA was reverse transcribed using Quantitech Reverse kit (Applied Biosystems). Quantitative PCR was performed with SYBR green Master Mix (Applied Biosystems) on an AB 7900-Fast-384 machine. Expression values were normalized to the expression of *Actb* and *Ywhaz* (whose expression as measured in our RNA-seq data did not change upon *Oct4* or *Nanog* induction) using the comparative *C*T method (54) and standard deviations were calculated and plotted using Prism 7.0 software (GraphPad). All assays were performed in triplicate.

### *In situ* hybridization

*In situ* hybridization in whole mount embryos or sections was performed using digoxigenin-labeled probes as described (55). Probes for *Hoxb1*, *Hoxb4*, *Hoxb9* and *Otx2* were generated by PCR, and the probe for *Oct4* was kindly provided by Tristan Rodriguez (Imperial College London). Early embryos were staged according to Forlani et al (2003).

### 4C-seq

4C was performed as previously described (32, 33) on two replicates of pools of 60-70 E9.5 embryos or 1-2×10^6^ G4 ES cells. Samples were crosslinked with 2% PFA, frozen in liquid nitrogen and stored at −80º. Chromatin was digested with *Dpn*II (New England BioLabs) followed by *Nla*III (New England BioLabs), and ligated with T4 DNA Ligase (Promega). For all experiments, 0.5-1 µg of the resulting 4C template was used for the subsequent PCR. 4C libraries were sequenced (single-end) at the CNIC Genomics Unit using an Illumina HiSeq 2500 sequencer. Sequences were mapped and quantified using RSEM v1.2.20 to the Mouse Genome Reference NCBIM37. Reads located in fragments flanked by two restriction sites of the same enzyme, in fragments smaller than 40 bp or within a window of 10 kb around the viewpoint were filtered out. Mapped reads were converted to reads per first enzyme fragment ends and smoothed using a 30-fragment mean running window algorithm. Smoothed scores from each experiment were then normalized to the total number or reads before the visualization. To calculate the frequency of captured sites per window Fastq files were demultiplexed using Cutadapt with the view point sequences as indexes. Potential Illumina adaptor contaminants and small chimeric reads were removed. Processed reads were assigned to their corresponding genomic fragment after a virtual digestion of the reference genome with the first and second restriction enzymes. Reads located in fragments 5kb around the viewpoint were filtered out. Quantification was performed considering each fragment end as one capture site if one or more sequences were mapped to it. The number of capture sites was summarized per 30 fragments window. The frequency of capture sites per window was used to fit a distance decreasing monotone function and Z-scores were calculated from its residuals using a modified version of FourCSeq (56). Significant contacts were considered in cases where the z-score was >2 in both replicates and deviated significantly (adjusted p value <0.05) from its normal cumulative distribution in at least one of the replicates.

### Chromatin immunoprecipitation

Chromatin immunoprecipitation (ChIP) was performed using 10 9.5 embryos or 1x 10^6^ ES cells per experiment. After recovery, embryos were treated with Collagenase Type I (Stemcell technologies; 07902) at 0,125% for 1h at 37º. Then, embryos were desegregated using a pipette and washed with cold PBS. Samples were fixed and protein-DNA complexes cross-linked by treatment with 1% formaldehyde (Pierce, 289069) for 15 min rocking at RT. To stop fixation, glycine (Nzytech, MBO1401)) was added to a final concentration of 125 mM during 10 min. Next, ChIP was performed using the ChIP-IT High Sensitivity kit (Active motif; 53040), following manufacturer’s instructions. DNA was sheared into fragments ranging from 200 bp to 1000 bp using a sonicator (Diagenode Bioruptor Water Bath Sonicator, 30 seconds ON 30 seconds OFF for 30 minutes). Immunoprecipitations was carried out using rabbit polyclonal anti-OCT4 antibody (Abcam; ab19857), and anti-Rabbit IgG polyclonal antibody (Abcam; ab171870) was used as negative control. Enrichment was measured by qPCR. A fragment from the *Nanog* promoter was used as a positive control (29, 31), and genomic fragments from the loci of *Anks1b*, *Smg6* and *Tiam1* as negative controls (57), after checking they did not contain OCT4 bound peaks (29).

### ES culture, cell editing and differentiation

ZHBTc4 cells were maintained in culture on 0.1% gelatin (sigma) using corning p24 plates with cell bind surface. Medium contained inactivated fetal calf serum (Hyclon), LIF (produced in-house) and 2i (PD0325901 and CHIR99021 Sigma). After one day in culture cells were treated with 1ug/ml tetracycline (Sigma) to turn of *Oct4*. Cells were collected from day 1 to 3 from the start of treatment. CRISPR/Cas9-mediated deletions in G4 mouse ES cells were generated using two guide RNAs together with a plasmid for Cas9. Cells were transfected, selected by sorting, and replated.

G4 ES cells (control), clone #30 and clone #57 were differentiated as described (45, 46) in monolayer using corning p24 plates with cell bind surface and with 0.1% gelatin (Sigma) added 30 min before passing. Cells were grown in N2B27 media supplemented with 10 ng/ml bFgf (R&D) for 3 days (d1–d3) and then were transferred into different media depending on the differentiation process. To induce hindbrain identity 10 nM RA (Sigma) was added from D3–D5. Spinal cord identity was induced by the addition of 5 µM CHIR99021 (Sigma) from D2 to D3 followed by 100 nM RA from D3–D5. To induce mesodermal differentiation the cells were treated with CHIR990215uM from D2–D5. Cells were collected at each time point by adding lysis buffer directly to the wells.

**Table S1.** Differentially expressed genes upon doxycycline treatment of *Oct4^tg^* or *Nanog^tg^* inducible models in E4.5 to E7.5 and E6.5 to E9.5 time windows.

**Table S2.** Clusters generated by unsupervised hierarchical clustering of genes differentially expressed in at least one condition.

**Table S3.** Gene Ontology analysis of clusters generated by unsupervised hierarchical clustering of genes differentially expressed in at least one condition.

**Table S4.** Primers and oligonucleotides used in this study.

## ACKNOWLEDGEMENTS

We wish to thank Beatriz Fernandez-Tresguerres for initial input to this work; Manuel Serrano and Konrad Hochedlinger for the *Nanog^tg^* mouse line; Austin Smith for the ZHBTc4 ES cell line; Tristan Rodriguez for reagents and discussions; Teresa Rayon, Miguel Torres and Moisés Mallo for comments and suggestions; Juan J. Tena for advice; the CNIC Genomics and Transgenesis Units for assistance; Simon Bartlett for English editing; and members of Manzanares lab for continued support. This work was supported by the Spanish government (grants BFU2014-54608-P and BFU2017-84914-P to MM; BFU2016-74961-P and BFU2016-81887-REDT to JLGS), the Andalusian Government (grant BIO-396 to JLGS) and the European Research Council (ERC) under the European Union’s Horizon 2020 research and innovation program (grant agreement No 740041 to JLGS). RR and RDA held FPU fellowships from the Spanish Ministry of Education, Culture and Sports (MECD). JV is a recipient of a “La Caixa” Fellowship and a Graduate Fellow of the Madrid City Council) at the Residencia de Estudiantes. Work in the lab of JLGS is supported by a María de Maetzu Unit of Excellence Grant (MDM-2016-0687) to the Department of Gene Regulation and Morphogenesis of the CABD. The CNIC is supported by the Spanish Ministry of Science, Innovation and Universities and the Pro CNIC Foundation, and is a Severo Ochoa Center of Excellence (SEV-2015-0505).

---

**Fig. S1.**
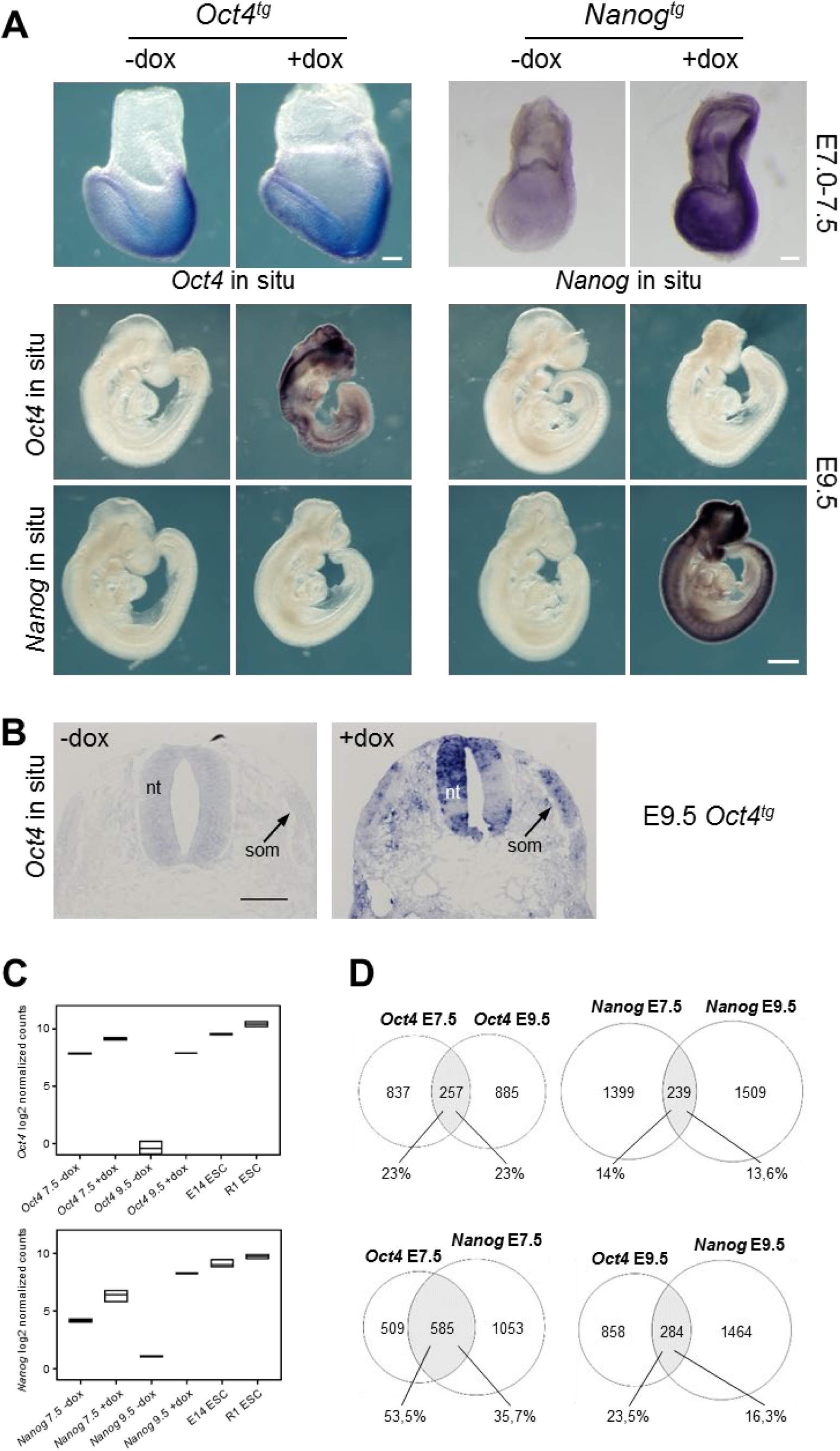
Characterization of the *Oct4^tg^* or *Nanog^tg^* mouse models. (*A*) Expression of *Oct4* and *Nanog* as determined by whole mount in situ hybridization in E7.0-7.5 and E95 untreated (controls, -dox) and treated (+dox) embryos from both transgenic lines. At E9.5, these factors do not cross-regulate. Scale bars, 100 µm (E7.0-7.5), 500 µm (E9.5) (*B*) In situ hybridization on sections of E9.5 untreated (controls, -dox) and treated (+dox) *Oct4^tg^* embryos showing induced *Oct4* expression in the neural tube (nt) and somites (som). Scale bar, 100 µm. (*C*) Comparison of the expression levels of *Oct4* and *Nanog* between *Oct4^tg^* or *Nanog^tg^* E7.5 and E9.5 embryos, without or with dox, and two strains of ES cells (E14 and R1), as log2 of normalized CPMs (counts per million) from RNA-seq data. (*D*) Overlap between genes differentially expressed in the *Oct4^tg^* or *Nanog^tg^* mouse models upon dox treatment in two 3-day time windows (up to E7.5 and up to E9.5).

**Fig. S2.**
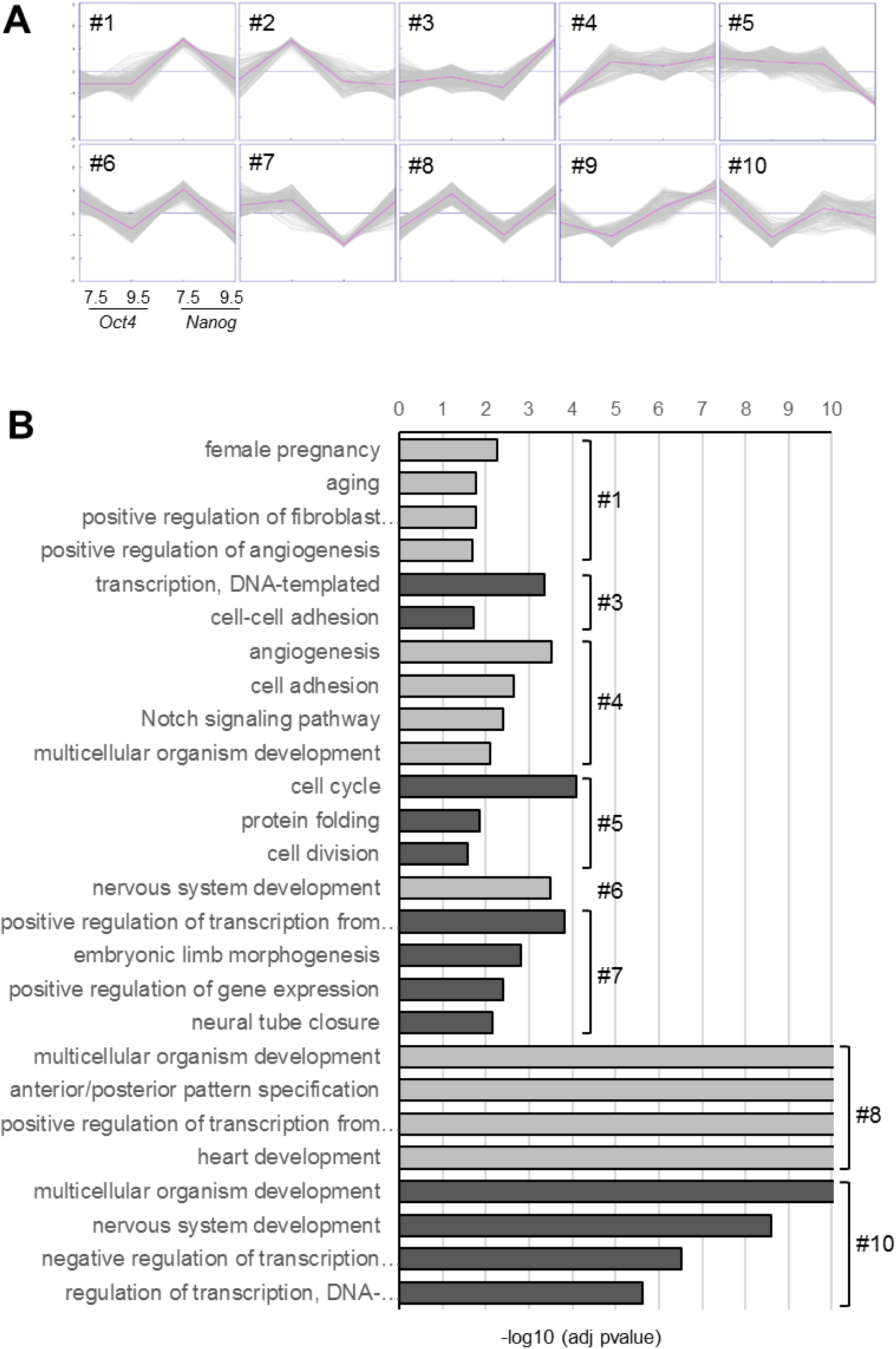
Transcriptomic analysis of changes induced by *Oct4* and *Nanog* expression in the mouse embryo. (*A*) Grouping of differentially expressed genes by k-means clustering, showing the response to *Oct4* or *Nanog* expression in the two time windows analyzed; y axis, z-scores. (*B*) Enriched categories for each cluster with a P value <0.05 (Bonferroni adjustment). Clusters #2 and #9 do not show significant enrichment for any category. Selected categories are shown; full listings are provided in Dataset S3.

**Fig. S3.**
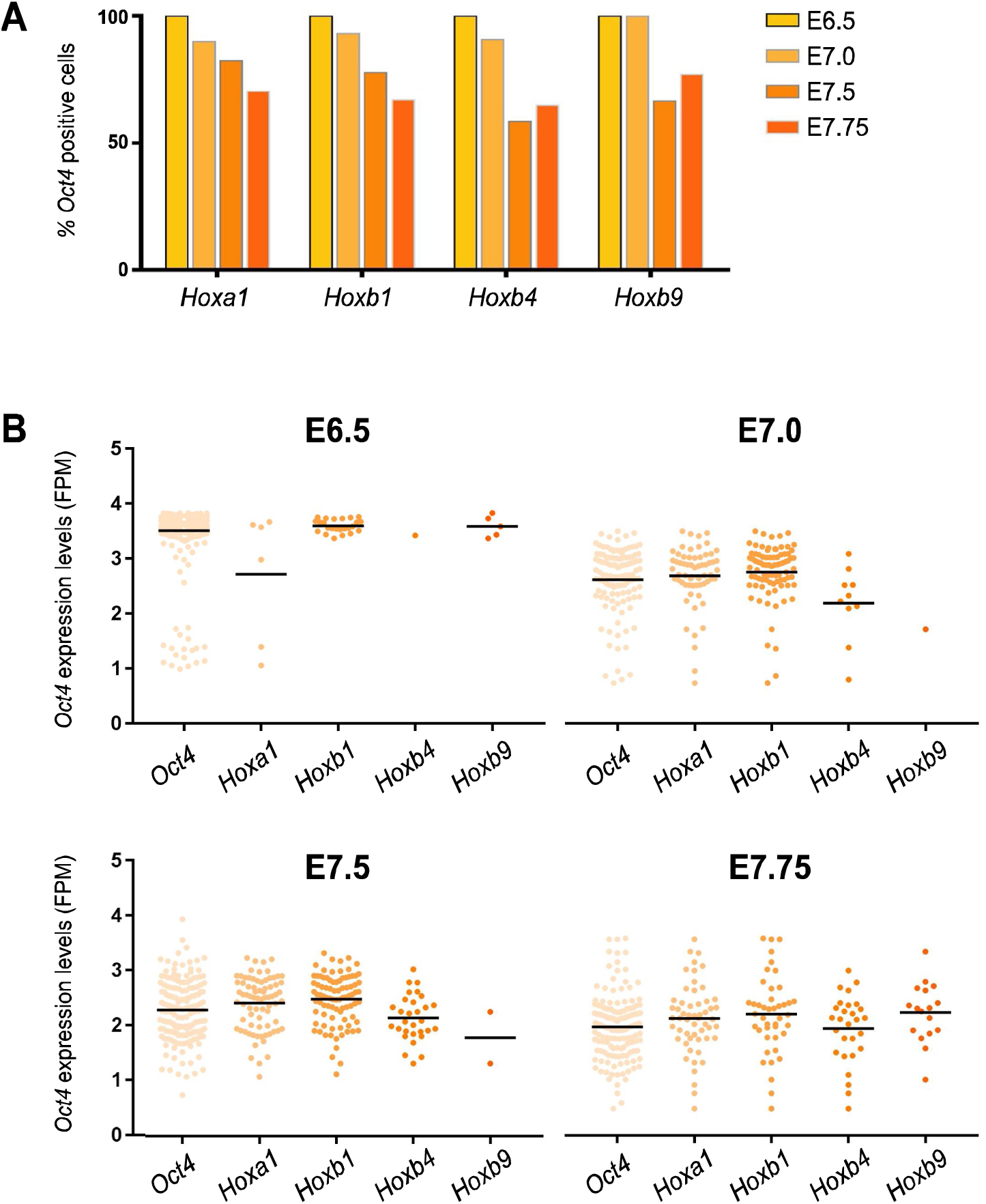
Co-expression of *Oct4* and Hox genes in the early postimplantation embryo. (*A*) Percentage of single cells expressing *Hoxa1*, *Hoxb1*, *Hoxb4* and *Hoxb9* at E6.5-E7.75 that also express *Oct4*. (*B*) Expression levels of *Oct4* in single cells co-expressing Hox genes at E6.5-E7.75, as log10 of FPM values; the rightmost column from each graph shows the expression levels of *Oct4* in all positive cells at that stage. Horizontal black bars indicate mean values. Single cell RNA-seq data from early embryos was obtained from Scialdone et al. 2016.

**Fig. S4.**
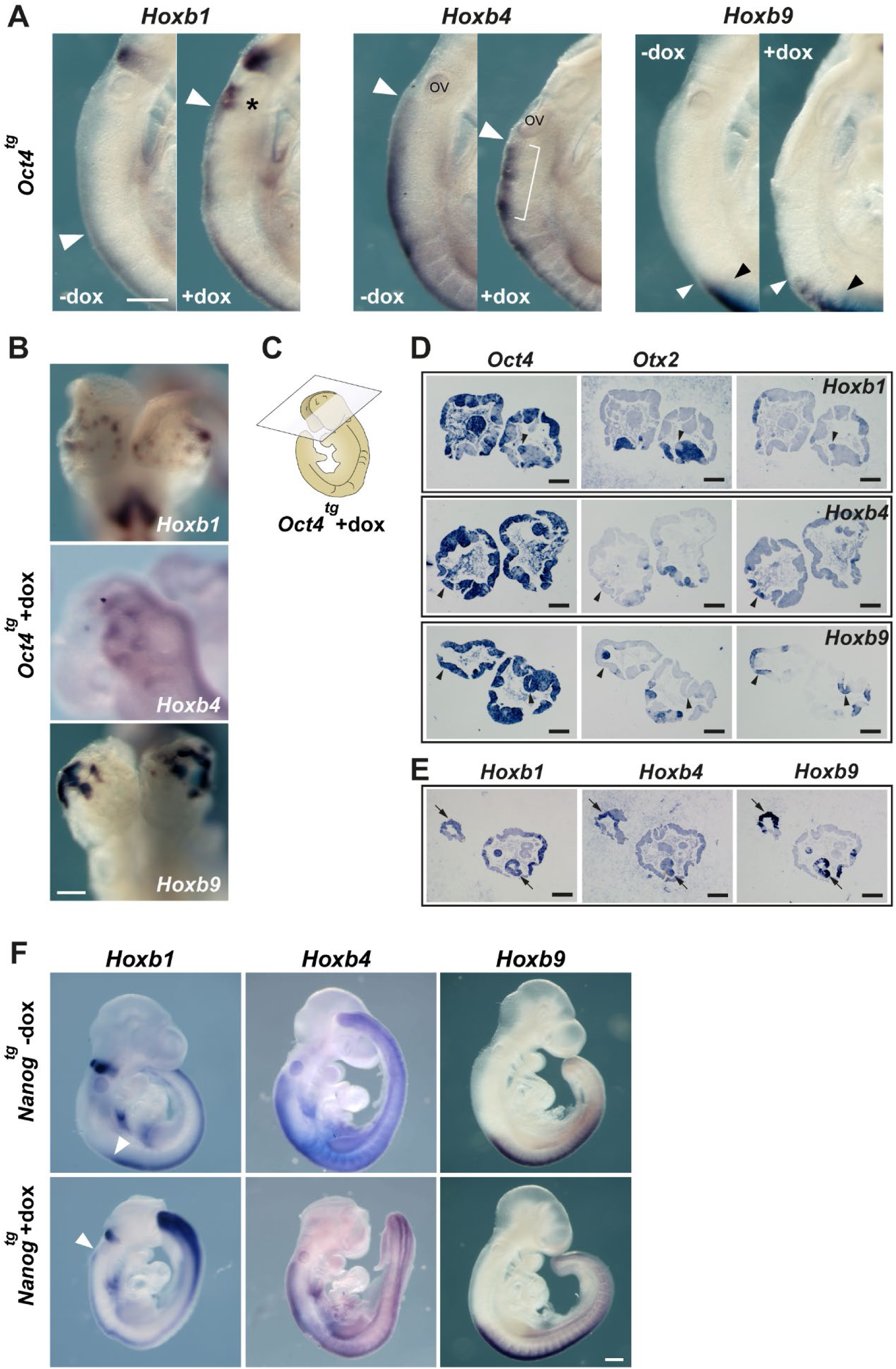
Changes to Hox gene expression patterns by pluripotency factors. (*A*) Close-up views of E9.5 embryos show in Fig. 2D. The expression of *Hoxb1* is expanded anteriorly (white arrowhead) in dox treated *Oct4^tg^* embryos and higher levels of expression are observed in presumptive rhombomere 6 (asterisk). Neural expression of *Hoxb4* shows a patchy and disorganized pattern upon dox treatment (bracket) but no change in its anterior limit (white arrowhead). Neural expression of *Hoxb9* is shifted anteriorly (white arrowhead) in relation to its limit of expression in the paraxial mesoderm (black arrowhead). Scale bar, 250 µm. (*B*) High-magnification anterior views of HoxB gene expression in vesicle-like structures in the anterior neural tube of dox-treated *Oct4^tg^* embryos. Scale bar, 250 µm. (*C*) Diagram indicating the section plane of dox-treated *Oct4^tg^* embryos shown in *D* and *E*. (*D*) In situ hybridization on consecutive sections for *Oct4* and *Otx2* together with *Hoxb1* (top), *Hoxb4* (middle) or *Hoxb9* (bottom); arrowheads indicate *Oct4* expression domains that have lost *Otx2* and gained Hox gene expression. Scale bars, 250 µm. (*E*) In situ hybridization on consecutive sections for *Hoxb1*, *Hoxb4* and *Hoxb9*, showing their co-expression (arrows). Scale bars, 250 µm. (*F*) Whole mount in situ hybridization of HoxB cluster gene expression in E9.5 controls (-dox, top row) and treated (+dox, bottom row) *Nanog^tg^* embryos. White arrowheads indicates the anterior limit of expression of *Hoxb1* in the neural tube. Scale bar, 250 µm.

**Fig. S5.**
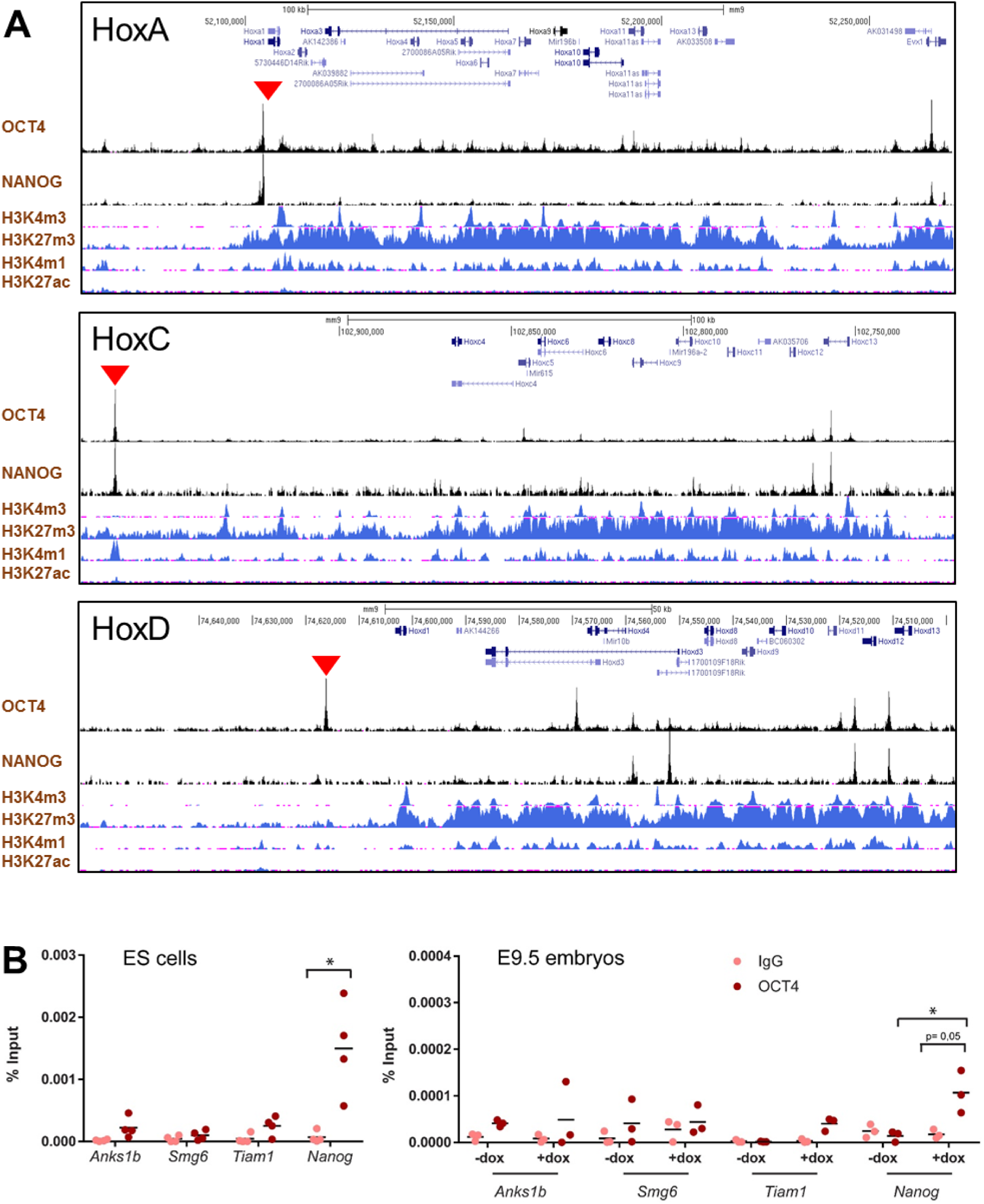
(*A*) ChIP-seq binding profiles for OCT4 and NANOG (29) along HoxA (mm9; chr6:52,059,532-52,270,099), HoxC (mm9; chr15:102,720,415-102,975,124; reversed) and HoxD (mm9; chr2:74,498,129-74,662,259; reversed) clusters in ES cells. Distribution of histone marks for active transcription (H3K4m3), repression (H3K27m3) and active regulatory elements (H3K4m1, H3K27ac) in Bruce4 ES cells (58) is shown below. Red arrowheads indicate strong binding of pluripotency factors at the anterior end of the clusters. (*B*) ChIP-qPCR of OCT4 binding to the positive (*Nanog* promoter) and negative (*Anks1b*, *Smg6* and *Tiam1*) control regions in ES cells (left panel, n=4) and in untreated (-dox) and treated (+dox) E9.5 *Oct4^tg^* embryos (right panel, n=3). Enrichment is shown as percentage of input for IgG (negative control) and anti-OCT4 antibody. Horizontal bars indicate means. * *P* value < 0.05 by two-tailed Student *t*-test.

**Fig. S6.**
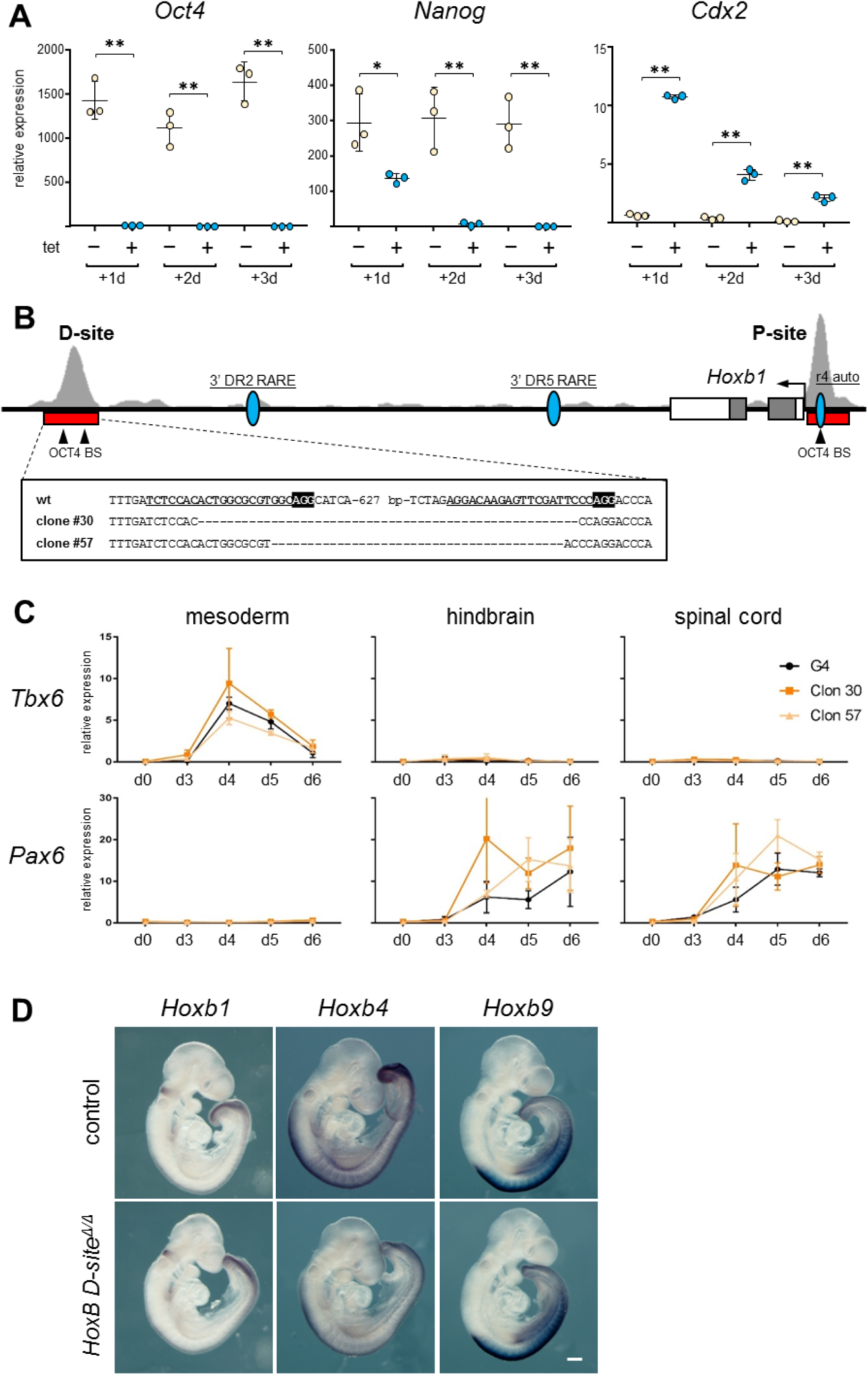
Characterization of *Oct4* depleted, and OCT4 D-site deleted ES cells and embryos. (*A*) Expression of *Oct4*, *Nanog* and *Cdx2* as measured by RT-qPCR in untreated (-, pale yellow) and tetracycline (tet) treated (+, blue) ZHBTc4 ES cells for three days (d1-d3), what causes *Oct4* downregulation. Statistical significance was calculated by Fisher’s Least Significant Difference test; *P* value: * <0.05, ** <0.01. (*B*) Genomic organization of the *Hoxb1* locus (mm9; chr11:96226282-96237333). Exons are indicated by boxes and the position of the *Hoxb1* promoter is indicated by an arrow. The location of distal (D) and proximal (P) binding sites is indicated by a red bar below ChIP-seq tracks for OCT4 (29) shown in light gray. The position of previously described regulatory elements for *Hoxb1* (the r4 auto-regulatory element, r4 auto; and two retinoic acid response elements, 3’ DR2 and 3’ DR5) is indicated by blue ovals. Arrowheads show the positon of consensus OCT4 binding sites located within the D and P sites. Below (boxed) the sequences of the deleted clones generated by CRISPR/Cas9 editing are shown. gRNAs are underlined and in bold type, and the PAM is indicated in black background and bold white lettering in the wild type (wt) sequence. (*C*) Expression as measured by RT-qPCR of a mesodermal (*Tbx6*) and a neural (*Pax6*) marker in control ES cells (G4, black line) and clones deleted for the distal OCT4 binding site (#30, dark orange; #57, light orange) during six days of differentiation along mesodermal, hindbrain, or spinal cord lineages. Expression is shown as relative levels. Error bars indicate SD, n=3. (*D*) Whole mount in situ hybridization for *Hoxb1*, *Hoxb4* and *Hoxb9* cluster in control (top row) and *HoxB D-site^Δ/Δ^* (bottom row) E9.5 embryos. Scale bar, 250 µm.

